# Optimization of Crop Rotation Calendar to Maximize System-Level Productivity under Climate Change

**DOI:** 10.1101/2025.05.25.656037

**Authors:** Keach Murakami, Toshichika Iizumi

## Abstract

Crop rotation offers multifaceted benefits for climate adaptation and mitigation. As rising temperatures accelerate crop development, it is necessary to adjust the rotation order and sowing dates of crops in a rotation system. Such an adjustment can enhance agricultural productivity at the whole rotation system level even under warmer conditions. Here, we present a computational approach to identify the crop rotation calendar (the order and sowing dates of rotation crops) that maximizes total crop output with available climatic resources. In this study, the rotation order indicates the cyclical sequence of crop species without intercropping, cover crop and fallow. This approach involves two steps: (1) generating crop windows (the timing and length of crop duration from sowing to harvest) and corresponding outputs (production per unit land area per harvest) for the rotation crops using a crop growth model with daily weather inputs; and (2) applying an optimization algorithm to identify a single subset of crop windows of rotation crops that maximizes the total crop output while satisfying agronomic constraints. We applied this method to a four-year, four-crop rotation system in northern Japan, with historical and projected climates from 2000 to 2100. The results revealed that adjusting crop windows would reduce the time to complete all rotation crops in the system (shortened rotation cycle) and enhance the system-level annual productivity [kgDW ha^−1^ year^−1^] with time. The increase in this productivity is approximately 0.5 % per year under high warming of the Shared Socioeconomic Pathway 5-8.5. Optimizing the rotation order in addition to the rotation crops sowing dates showed the changes in the most productive rotation orders, demonstrating the dynamic nature of rotation planning under climate change. Our methodology offers a flexible, scalable means of designing cropping system to harness the positive effect of climate change on long-term food security.

## Introduction

Crop rotation is defined as an agricultural practice of cultivating different plant species on the same land with a specific rotation order (Fig. 1). This practice is adopted mainly because of its yield-related benefits arising from disrupting cycles of weeds, pests and pathogens and from improving soil health by cultivating different crops in succession (Bullock, 1992). A research synthesis of long-term trials at sites in Europe and North America found that diversifying crop functional types in a rotation system enhanced yields of maize and small grain cereals (Smith et al., 2023). Furthermore, studies suggest that crop rotation serves as an effective adaptation measure to climate change. Using the same dataset as Smith et al. (2023), Costa et al. (2024) reported that diversified crop functional types in a rotation compensated yield losses from anomalous warm conditions, long and warm dry spells, as well as from anomalous wet conditions for small grains and dry conditions for maize. Marini et al. (2020) reported that crop rotation mitigated cereal yield losses caused by adverse climates, such as drought and anomalous temperatures, compared to monoculture. Moreover, crop rotation is reported to reduce greenhouse gas emission from agricultural fields (Saghai et al., 2025; Yang et al., 2024), underscoring the significance of this practice for climate mitigation.

**Fig. 1.**
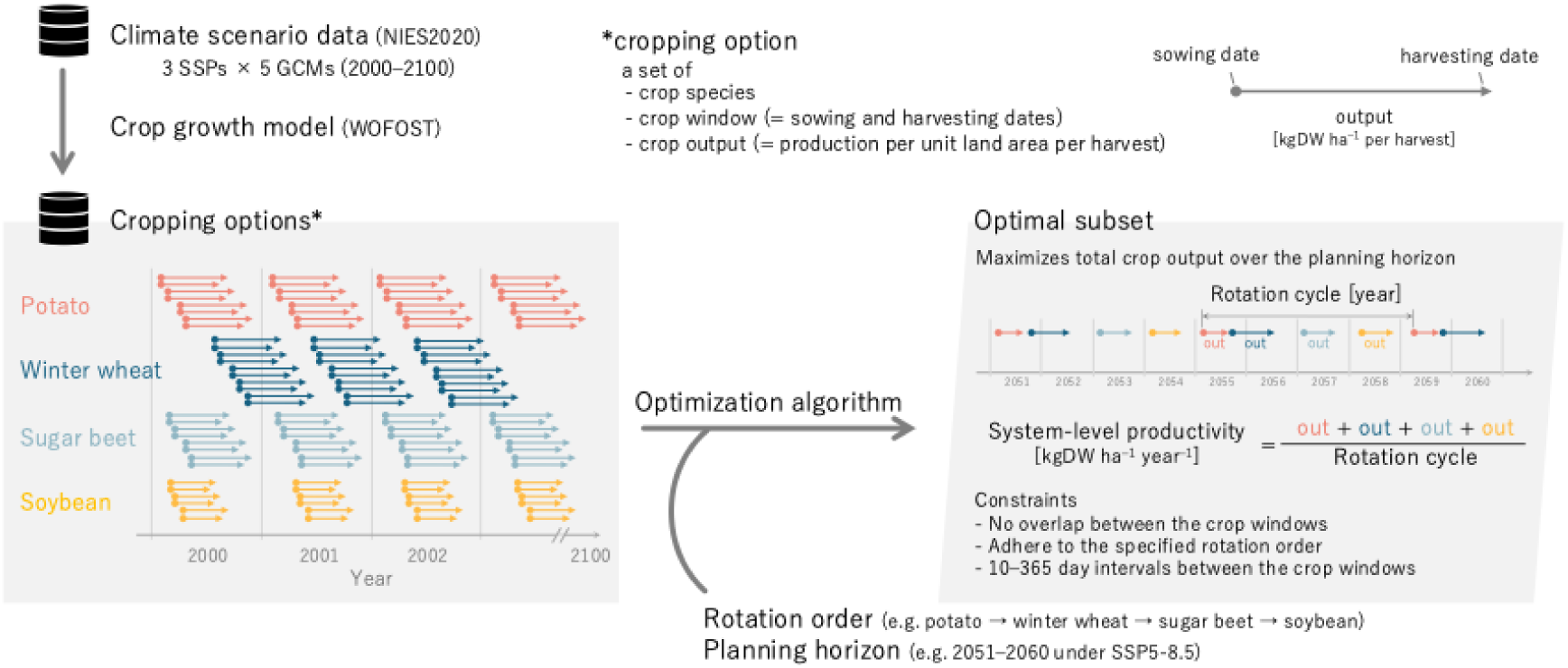
Diagram of optimization of crop rotation calendar proposed and definitions of concepts in this article. A large number of cropping options are generated by using a crop growth model driven by climate scenario data. The optimization algorithm identifies a subset of the cropping options that maximize total output of rotation crops that satisfies agronomic constraints.

Climate adaptation requires adjustments in the crop window—the timing and length of crop duration from sowing to harvest—of rotation crops (hereafter referred to as the crop rotation calendar). This is because warmer climates accelerate crop development and therefore shorten the length of crop duration. In extreme cases, the time required to complete a two-crop sequence can shorten from two years under current climate to a single year in warmer conditions. Such an adjustment in the crop rotation calendar may reduce the rotation cycle—the time required to complete a cycle of rotation—and enhance agricultural productivity at a whole rotation system level. If two crops were harvested in a year, then the system-level annual productivity doubles. Therefore, adjusting the crop rotation calendars is crucial for maximizing the positive effect of climate change on agricultural productivity for efficient land use.

Shifting sowing date of a single crop has been intensively studied as an adaptation measure (e.g. Asseng et al., 2013). However, optimizing the crop rotation calendar remains underexplored. For example, to optimize the rice-wheat rotation system in the Indo-Gangetic plain, Urfels et al. (2022) found that shifting rice sowing dates affected not only rice yields but also the subsequent wheat yields. Similarly, by shifting sowing dates of both crops by ±30 days in rice-wheat rotation and rice-rice double cropping systems, Wang et al. (2022) sought optimal sowing dates in response to climate change in India and Bangladesh. They used single-objective optimization to maximize annual caloric yield and multi-objective optimization to minimize blue water requirements with minimal yield reduction using a genetic algorithm. Such a date-shifting approach becomes complex depending on the site-specific climatic constraints and the number and functional type of rotation crops in a system. Adjusting the window of one crop may affect the preceding and following crops, potentially disrupting the entire rotation. A cycle of crop rotation sometimes contains seasonal gap periods, such as long winters in high latitudes or dry seasons in rainfed systems. For successful harvesting of winter crops, young seedlings should be exposed to a certain level of low temperatures to complete vernalization. The number of crop windows and constraints that needs to be considered in the date-shifting approach would become increasingly complex, posing a computational challenge.

Optimizing the crop rotation calendar has been formulated as a seasonal crop selection problem and often solved using linear programming techniques (Dury et al., 2012; El-Nazer & McCarl, 1986; Klein Haneveld & Stegeman, 2005; Detlefsen & Jensen, 2007; Salassi et al., 2013; Tsai et al., 1987; Santos et al., 2011). Several studies have adopted dynamic programming as a more efficient method. Burt & Allison (1963) used dynamic programming focusing on soil moisture to support farmers in selecting either wheat cultivation (water consumption) or fallowing (water replenishment) for given seasons. Taylor & Rodrıguez-Kábana (1999) modeled population dynamics of two peanut pests and used dynamic programming to identify optimal patterns for peanut-cotton rotation in the southeastern United States. Trengove & Manson (2003) proposed a profit-maximizing rotation order through dynamic programming based on crop yields calculated from water availability, yield penalties from monoculture, and production costs. By combining this method with assumptions on changes in growing-season rainfall, they simulated climate change effects on gross margins. However, these studies only considered a single sowing date for each crop and season. To the best of our knowledge, no study has considered flexible sowing dates of rotation crops in conjunction with their rotation order in a changing climate.

Here, we present a methodology for optimizing the crop rotation calendar that maximizes system-level annual productivity of crop rotation under a given climatic condition (Fig. 1). In this method, a large number of ‘cropping options’ of rotation crops are generated by a crop growth model. A cropping option is a set of crop window (the timing and length of crop duration from sowing and harvest) and crop output (production per unit land area per harvest) of a crop species. Because the term ‘yield’ usually refers to one harvest per year implicitly, we here use output instead. Daily data from the projected climates are used to drive the crop growth model. The proposed method then identifies a subset of cropping options that maximizes total crop output using an optimization algorithm based on dynamic programming. By applying this method for historical and projected climates, we investigate the optimal crop rotation calendar and their future changes. A four-crop, four-year rotation system popular in Hokkaido, the northern part of Japan is selected as a testbed. This region is characterized by a combination of summer crops, such as sugar beet, potato, and soybean, with winter wheat that overwinters under seasonal snowpack.

## Materials and Methods

### Dataset

We generated cropping options by running the WOFOST crop growth model (de Wit et al., 2019) forced with the NIES2020 meteorological forcing dataset (Ishizaki, 2021). The dataset of cropping options is available in Supporting information Data S1. The analysis focused on a representative location (42.9°N, 143.1°E) located within an upland farming region in Japan’s northernmost island, Hokkaido. This site is characterized by seasonal snowpack from November to April and sufficient water supply during the growing season.

We used daily data on minimum and maximum temperatures, precipitation, and solar radiation for the historical period from 2000 to 2014, followed by the future period from 2015 to 2100. Outputs of five CMIP6 general circulation models (GCMs), including ACCESS-CM2, IPSL-CM6A-LR, MIROC6, MPI-ESM1-2-HR, and MRI-ESM2-0, under three Shared Socioeconomic Pathways (SSPs; SSP1-2.6, SSP2-4.5, and SSP5-8.5) were used. Snow water equivalent was calculated from minimum and maximum temperatures and precipitation following Trnka et al. (2010) and regarded as the indicator of snow cover. CO_2_ concentration scenarios in CMIP6 were used as an input of the crop growth model (Riahi et al., 2017).

We used the R software (R Core Team, 2023) and its implementation of WOFOST (Hijmans, 2021) to calculate potential yields under a given crop window and growing-season climate. This approach assumes no yield reductions from diseases and pests, weeds, nutrient deficiencies, or water stress. Crop parameters for the four species were selected from default parameter sets in Rwofost based on their ability to reproduce actual crop phenology under historical conditions. Specifically, we utilized wofost_crop(“wheat_spa”) for winter wheat, wofost_crop(“potato_704”) for potato, wofost_crop(“sugarbeet_604”) for sugar beet, and wofost_crop(“soybean”) for soybean. For each sowing date that is potentially feasible due to no snow cover, we simulated the temporal evolution of dry weight of storage organs (WSO) and developmental stage (DVS). For potato, if DVS was greater than 95 % of its maximum value, we considered the crop harvestable. The other three crops were considered harvestable if WSO was greater than 90 % of its maximum value. This approach thus allowed multiple potential harvesting dates for each sowing date. For winter wheat, we incorporated vernalization and overwintering constraints to remove invalid sowing dates (Murakami et al., 2024). To ensure vernalization for successful reproduction, the number of days with mean temperatures below 5°C was required to exceed 65 days before reaching a DVS of 0.45 (approximately young panicle formation). To ensure overwintering survival, growing degree days before the first day of snow cover was greater than 300 °C d. By iterating these simulations across all snow-free dates, we constructed a dataset of cropping options—simulated sowing and harvesting dates and outputs [kgDW ha^−1^ per harvest] for all rotation crops. This dataset was used for the crop rotation calendar optimization described in the following section.

### Optimization algorithm

We propose an algorithm to identify a subset of cropping options that maximizes the total output of rotation crops over a planning horizon while meeting three constraints (Fig. 1, Script Supporting information): (1) no temporal overlap between crop windows, (2) adherence to the specified rotation order, and (3) 10–365 day intervals between harvesting and subsequent sowing for practical feasibility. This algorithm extends the weighted non-overlapping interval scheduling, a typical design pattern of dynamic programming. For each of the five GCMs and starting years from 2000 to 2090, we identified an optimal subset of cropping options over a 10-year planning horizon. To simulate patterns initiated with four crops (winter wheat, sugar beet, soybean, or potato), the initial cropping option for each optimization was fixed as the highest-yielding one for the initial crop in the starting year. To eliminate dependency on this initial selection, the first four options—equivalent to one complete crop rotation—were regarded as spin-up and were not used for analyses. The rotation cycle was defined as the duration between sowing dates of the initial crop in the second and third rotations, rounded to the nearest integer in a unit of year. System-level annual productivity [kgDW ha^−1^ year^−1^] was defined as a sum of outputs of four crops in the second rotation divided by the rotation cycle. We evaluated the system-level annual productivity under the current rotation order (winter wheat-sugar beet-soybean-potato; WBSP) and other possible rotation orders (WBPS, WPBS, WPSB, WSBP, WSPB) (Supporting information Data S2). Under several planning horizons, crop rotation was not available because no subset satisfied the constraints. These cases were removed when summary statistics and trends were calculated. This removal had negligible effects because these cases hardly occurred.

To quantify the effect of the optimization, we also calculated system-level annual productivity under the projected climates with the current rotation order without optimization (WSBP in four years) (Supporting information Data S3). In this case, we extracted cropping options where the harvests of four crops occur in four serial years. From these cropping options, the system-level annual productivity was calculated from the subset that meets the three constraints. Four-year rotation was not available in some situations particularly because the interval between potato harvesting and winter wheat sowing was smaller than 10 days. These cases were removed from the calculation of summary statistics and trends.

## Results

### Cropping options, rotation cycle and crop rotation calendar

The effect of warming on the number of available cropping options varied depending on the crop species and warming levels (Supporting information Table S1). Potato and sugar beet, which require higher growing degree-days for maturation, showed an increase in the number of cropping options under warmer conditions. In contrast, soybean and winter wheat showed minimal changes or even decreases in some cases, in particular around 2100 under high warming scenario (SSP5-8.5). This pattern resulted from accelerated development due to warming, which shortened the number of days from harvestable condition to full maturity.

The rotation cycles identified by the optimization algorithm varied according to warming levels, but consistently showed a shortening trend with time (Fig. 2). Under SSP2-4.5 and SSP5-8.5, the frequency of three-year cycles exceeded 50 % around 2070 and 2050, respectively. Two-year cycles emerged in rare instances in SSP5-8.5 by the end of this century. Five-year cycles hardly occurred after 2070 and 2050 under SSP2-4.5 and SSP5-8.5, respectively. For SSP1-2.6, three-year cycles remained limited to approximately 10 % frequency and five-year cycles still occurred occasionally at the end of the century.

**Fig. 2:**
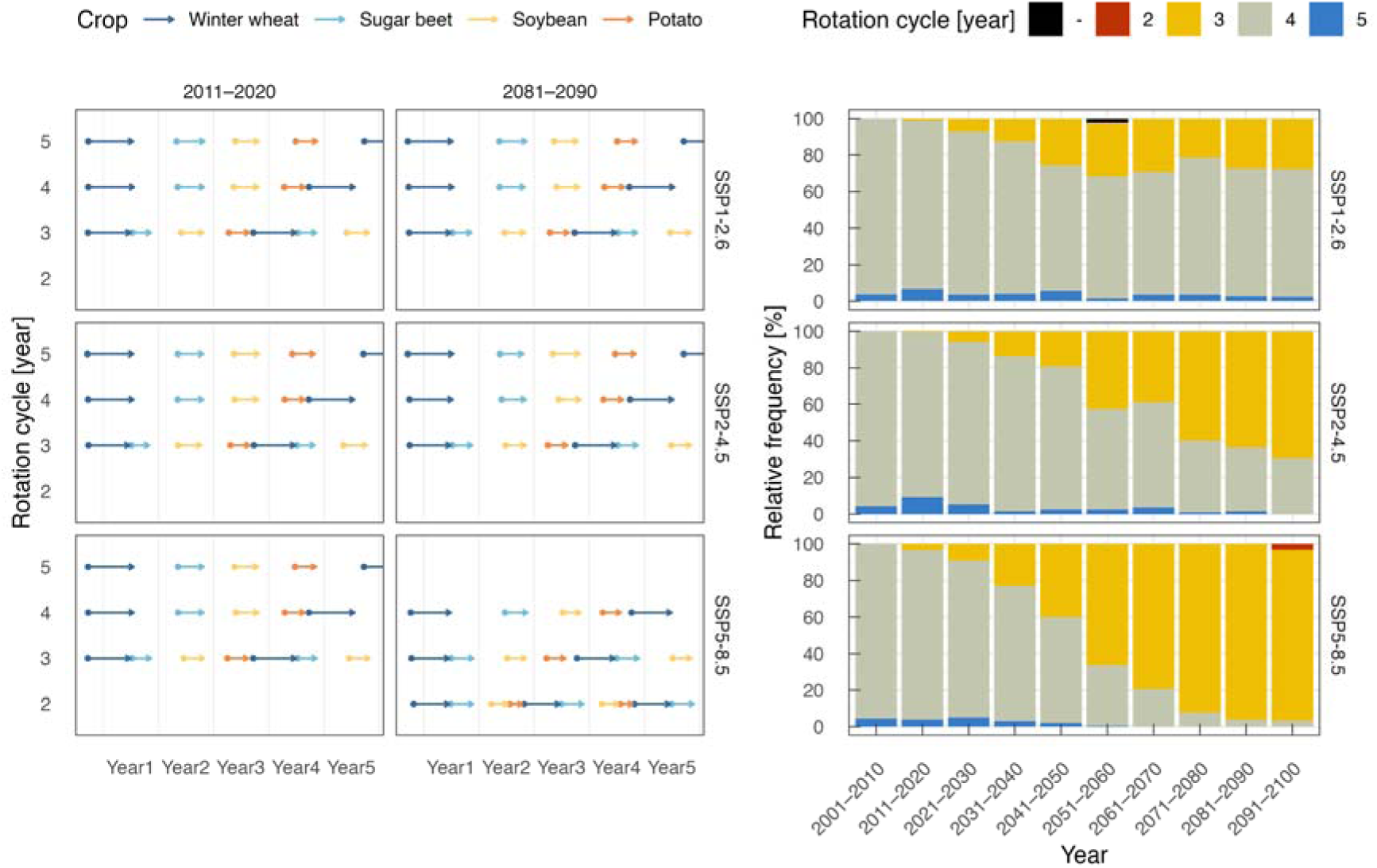
Crop calendars of four rotation crops identified by the optimization algorithm (left) and frequencies of crop rotation cycles (right) for winter wheat-sugar beet-soybean-potato rotation under different SSP scenarios and time periods. The start and end points of arrows indicate mean sowing and harvesting dates, respectively. Rotation cycles are not defined in instances where crop rotation is not available and shown in black in the right panel.

The crop rotation calendars—i.e., calendars of four rotation crops—identified by the optimization algorithm depended on the rotation cycle, but not on warming levels (Fig. 2). In five-year cycles, winter wheat sowing was postponed to the next year due to delayed potato harvesting compared to conventional four-year cycles. In three-year cycles, sugar beet was sown and harvested within the same year of winter wheat harvest. In the rare two-year cycles that became feasible under extremely warm conditions, winter wheat was sown in the same year after soybean and potato cultivation. As warming progressed, sowing and harvesting dates of winter wheat tended to shift later and earlier, respectively (Supporting information Fig. S1). Both sowing and harvesting dates shifted later for sugar beet and soybean under warmer conditions, while both dates shifted earlier for potato.

### System-level annual productivity and outputs of four rotation crops

System-level annual productivity (i.e. total output of rotation crops per unit land area per year) showed increasing trends in all scenarios with time (Fig. 3). Linear regression showed increases in system-level annual productivity of 5.84, 19.4, and 37.7 kgDW ha^-1^ year^-1^ under SSP1-2.6, SSP2-4.5, and SSP5-8.5, respectively. These increases were associated with the increased frequency of three-year rotation cycles and decreased frequency of five-year cycles. Compared to the conventional four-year crop rotation without optimization, system-level annual productivity estimated by the optimization algorithm was higher particularly in warmer conditions under high-emission scenarios and later in this century. This difference represents the benefit of optimization of crop rotation. Note that trends calculated from the non-optimized simulations removed the situations where four-year rotations were impossible. Because this systematically overestimated productivity of non-optimized simulations, the benefit of crop rotation optimization was even greater.

**Fig. 3:**
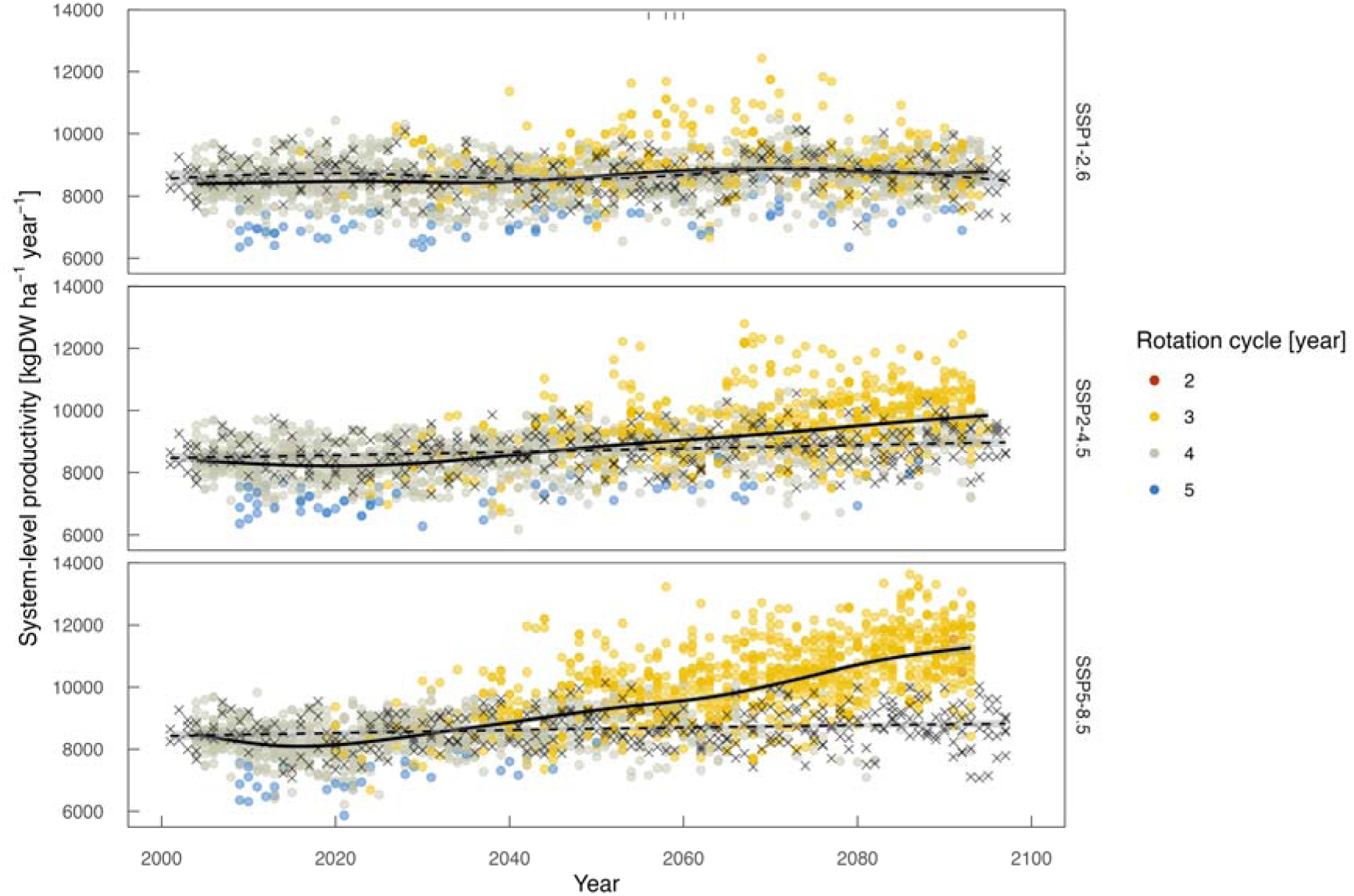
System-level annual productivity maximized by the optimization algorithm for winter wheat-sugar beet-soybean-potato rotation under different SSP scenarios and time periods. Values (circles) and trends (solid lines) are shown. Tick marks at the top of panels indicate instances where crop rotation is not available. Dashed lines and crosses represent system-level annual productivity without optimization where rotation cycles are fixed at four years.

Among the four crops, output of sugar beet was most strongly affected by optimization (Fig. 4). The output decreased substantially when rotation cycles shortened to three years. The sugar beet output in three-year cycles increased with time. Soybean and wheat outputs were not dramatically affected, but demonstrated increasing trends parallel to warming progression in SSP2-4.5 and SSP5-8.5 scenarios. Annual rates of increasing in output for soybean were 6.89, 12.5, and 15.0 kgDW ha^-1^ under SSP1-2.6, SSP2-4.5, and SSP5-8.5, respectively. For wheat, the annual rates of increasing were 3.26, 7.18, and 11.4 kgDW ha^-1^, respectively. For potato, although the trends were less obvious due to large interannual variations, the annual rates of increasing were 9.29 and 5.77 kgDW ha^-1^ under SSP2-4.5 and SSP5-8.5, respectively. SSP1-2.6 showed no statistically significant trend (*P* = 0.553).

**Fig. 4:**
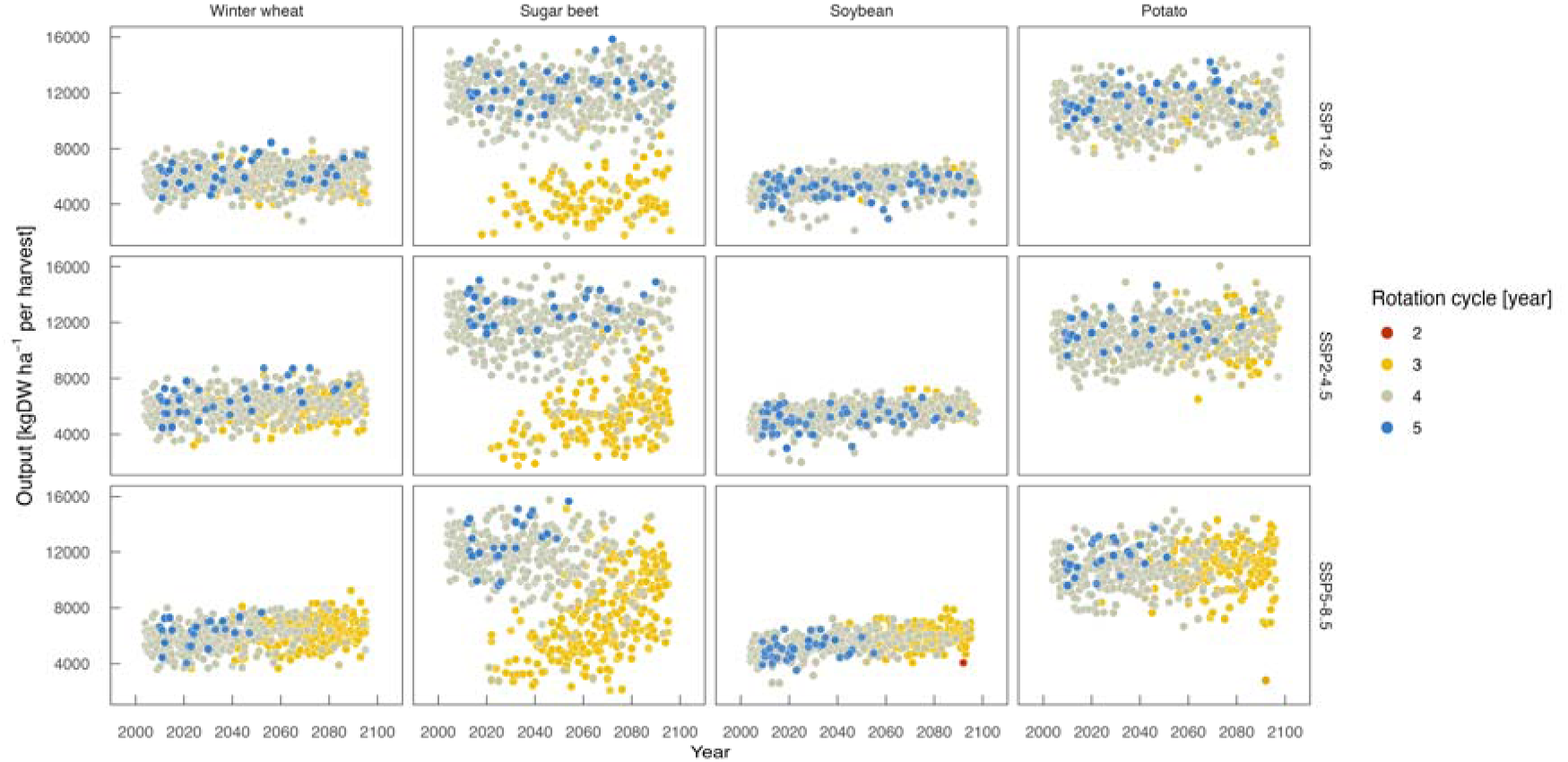
Outputs of four rotation crops identified by the optimization algorithm for winter wheat-sugar beet-soybean-potato rotation under different SSP scenarios and time periods.

### Determination of optimal rotation order

Under the historical climate condition around 2000–2020, system-level annual productivity was higher in WBSP (winter wheat-sugar beet-soybean-potato), a widely applied rotation order, and WSBP (winter wheat-soybean-sugar beet-potato) than in the other rotation orders (Fig. 5). WSBP exhibited substantial increase in system-level annual productivity with time in all scenarios. WSBP consistently showed the highest system-level annual productivity irrespective of scenarios and time periods. The increase in WBSP was slower than those in the other rotation orders and stagnated under SSP1-2.6. The four rotation orders with lower system-level annual productivity under the historical condition (WBPS, WPBS, WPSB, and WSPB) showed steady increases with time. Consequently, system-level annual productivity of these four rotation orders reached levels comparable to WBSP around 2040 irrespective of scenarios.

**Fig. 5:**
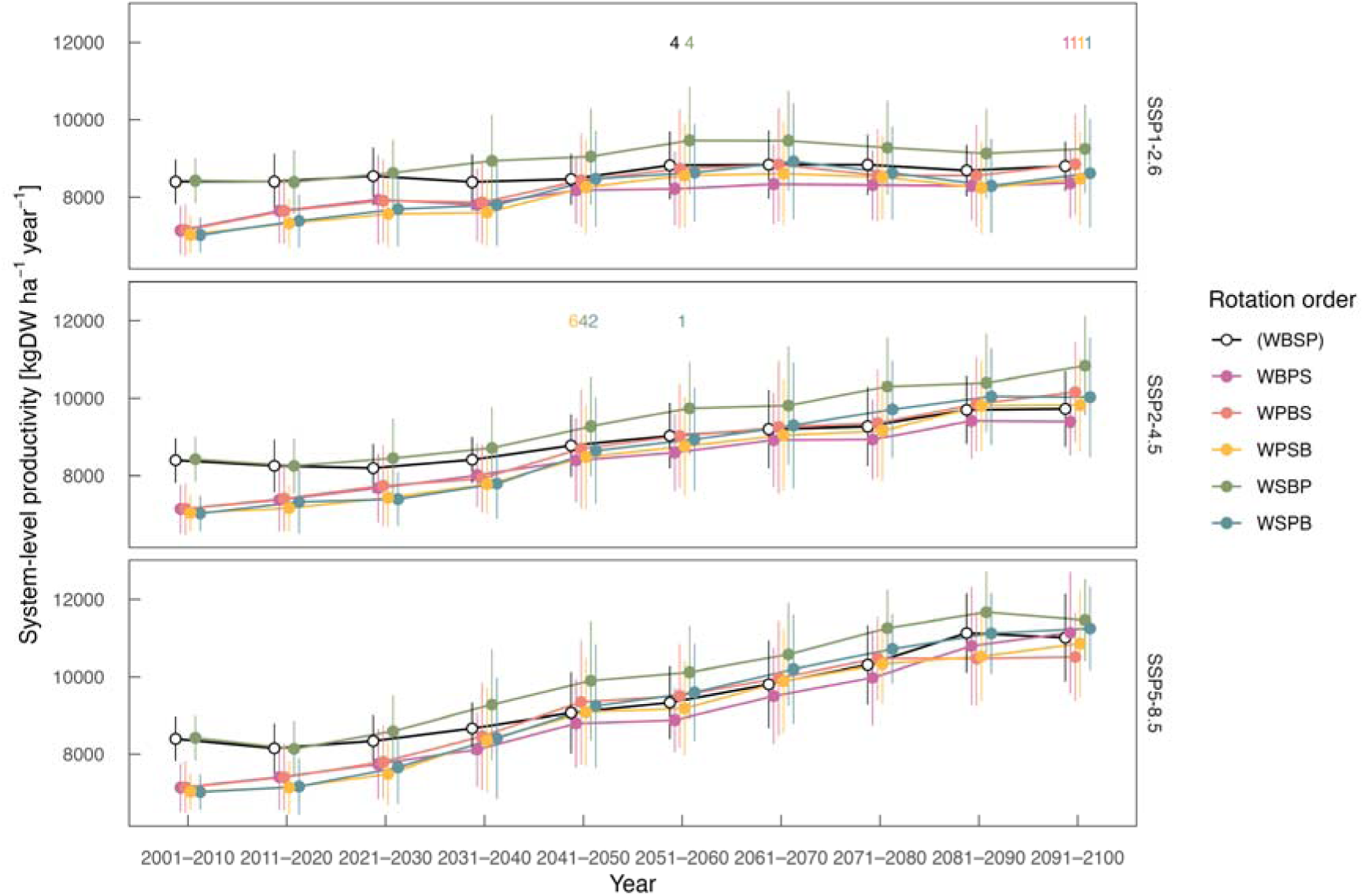
System-level annual productivity maximized by the optimization algorithm with different rotation orders under different SSP scenarios and time periods. Means (circles) and standard deviations (errorbars) are shown. Letters above the errorbars represent the number of instances where crop rotation is not available. The rotation order shown with parentheses represents the conventional rotation order. W: winter wheat, B: sugar beet, S: soybean, P: potato.

## Discussion

### Speeding up rotation cycle to boost productivity of crop rotation

This study demonstrated that reconstructing crop rotation calendars can increase the system-level total crop output per unit land area per year (or system-level annual productivity). While global cropland relocation (i.e. spatial optimization) has been studied to enhance land-use efficiency (Beyer et al., 2022), crop rotation is not incorporated. Considering the stagnation in crop productivity despite population growth (Gerber et al., 2024), optimizing crop rotation could serve as one solution to address global food security. Therefore, our study presents an additional pathway for enhancing crop productivity. The annual rate of increasing in system-level annual productivity calculated by simple linear regression under the SSP5-8.5 scenario was 37.7 kgDW ha^-1^ year^-1^, corresponding to a rate of approximately 0.5 % since system-level annual productivity was ∼8000 kgDW ha^-1^ year^-1^ around 2020 (Fig. 3). This contribution of crop rotation optimization to overall crop productivity is sizable compared to the required annual growth rate of 2.4 % and the observed rate of 0.9–1.6 % (Ray et al., 2013).

Warming accelerated crop development and shortened rotation cycles; This enhanced system-level annual productivity of crop rotation (Figs. 2, 3). The shortening of rotation cycles depends strongly on the rotation order. For example, in the conventional rotation order (winter wheat-sugar beet-soybean-potato) adopted in the present testbed, shortening of the rotation cycle occurred when sugar beet is sown and harvested in the same year after winter wheat harvest (Fig. 2). Substituting early-maturing varieties may enable earlier shortening of rotation cycles and enhance system-level annual productivity. In practice, a conventional rotation order is designed considering various factors including workload distribution and nutrient and water requirements of each crop. There should be substantial inertia against changing the established rotation practices. In regions where highly environment-adapted agriculture has developed and spatially uniform farming methods are implemented, changes in agricultural practices tend to occur less readily (Kling et al., 2024). Field trials that directly compare rotation orders are convincing but require considerable time. The present simulations help redesign crop rotation calendars for timely adaptation for climate change.

### Aspects to be considered for decision making in practice

The stability of crop rotation calendar is key for adoption by farmers, as well as productivity. During transitional periods when rotation cycle shortening may or may not occur, there is potential for crop failure. For example, shortening of the rotation cycle may not occur once in three times around 2050 even under the SSP5-8.5 scenario in the present testbed (Fig. 2). When farmers sow crops anticipating a shortened rotation cycle but the crops do not reach harvestable conditions, associated costs become losses. Long-term weather forecasts could mitigate such risk. Since temperature largely determines whether crops reach harvestable conditions, accurate long-term weather forecasts spanning several months would help avoiding crop failure. A recent study showed pre-sowing predictions using 8-month weather forecasts (Anderson et al., 2024, Iizumi et al., 2024).

Farmers may prefer to cultivate an additional crop during fallow periods resulting from optimization rather than drastically changing their rotation systems. For example, in the optimized three-year cycle crop rotation calendar (Fig. 2), there is a fallow period before soybean seeding. The present method could help evaluate the feasibility of introducing new crop species and varieties given that crop growth models are available. Our approach could also be used for crop calendar optimization in multiple cropping systems, which account for 34 %, 13 % and 10 % of the global rice, wheat and maize area, respectively (Iizumi & Ramankutty, 2015, Waha et al. 2020).

The accuracy and uncertainty associated with crop growth models should be addressed in future work. The present study used a single crop growth model with default parameters as a proof of concept. In multi-site wheat yield simulations, uncertainty attributable to crop growth models exceeded that from climate models (Asseng et al., 2013). It would be preferable to integrate crop and climate models within frameworks like AgMIP for individual crops (e.g. Jägermeyr et al., 2021). Then, system-level annual productivity should be optimized based on crop-level results in a hierarchical manner. Most climate change simulations have fixed sowing dates according to current practices (Jägermeyr et al., 2021, Müller et al., 2024) or shifted them by certain days (Asseng et al., 2013), implying that the identification of crop calendar transitions remains to be resolved. A key advantage of our method is that it automatically identifies sowing dates of rotation crops. This eliminates the need for arbitrary crop calendar settings in future projections.

The present approach does not account for positive effects of crop rotation on soil fertility (Bullock, 1992) since it assumes no nutrient deficiencies. Incorporating the positive feedback from soil fertility to crop yields (Henryson et al., 2018) may facilitate discussion on efficient fertilizer use for sustainable agriculture.

### Algorithm implementation and extensions

Interval scheduling, which forms the basis of the present optimization algorithm, is a typical application of dynamic programming (Bellman, 1966). A naive brute-force approach that evaluates all possible combinations of *N* cropping options (2*^N^* patterns) is computationally infeasible. Linear programming can reduce this cost to polynomial complexity. Dynamic programming further improves computational efficiency by exploiting the sequential structure of the problem. This advantage becomes particularly significant in large-*N* problems like this study where numerous cropping options exist over the planning horizon (Supporting information Table S1).

Our current implementation aims to maximize the dry weight of storage organs (i.e. grain, tuber, and root); however, the method can be applied to other target variables. If the objective is to maximize farmers’ profits, output values could be converted to revenue using crop-specific prices and production costs. Alternatively, to evaluate food supply potential for a growing population, dry weight can be converted to caloric output using appropriate crop-specific coefficients. A more realistic and critical challenge would be multi-objective optimization, such as simultaneously maximizing yield and minimizing water use (e.g. Wang et al., 2022). Multi-objective optimization exponentially expands the search space, making it difficult to identify the Pareto front. Heuristic methods such as genetic algorithms are likely necessary. For example, Jain et al. (2021) applied a hybrid approach combining the crow search algorithm and particle swarm optimization to optimize spatiotemporal crop patterns in India, aiming to increase net crop benefits while reducing fertilizer use.

## Conclusions

A computational approach for optimizing crop rotation calendars was developed to maximize system-level annual productivity under climate change scenarios. In this approach, an optimal subset of cropping options was identified from a large number of cropping options generated using a crop growth model driven by climate scenarios. This methodology demonstrated that optimizing rotation orders and sowing dates of rotation crops enhanced system-level annual productivity without varietal replacements. Crop rotation optimization represents an effective adaptation strategy, with expected shortened rotation cycle under warming. The present method is flexible and scalable when testing whether new varieties with different temperature requirements or novel crops would fit to a given rotation system. These findings provide practical guidance for cropping system design, including breeding programs and cultivation management strategies, to harness the positive effect of climate change on crop production.

## Acknowledge

K.M. was supported by JSPS KAKENHI 23K14051. T.I. was supported by the Environment Research and Technology Development Fund (JPMEERF23S21120) of the Environmental Restoration and Conservation Agency provided by the Ministry of the Environment of Japan.

## Data availability statement

The data and script that support the findings of this study are available as supplemental files in this paper (https://doi.org/XXXX). NIES2020 meteorological forcing data (ver. 1.2) are available at https://doi.org/10.17595/20210501.001.

## Supporting information

Supporting Information Script S1

## Notes

### Competing Interest Statement

The authors have declared no competing interest.

