## Supporting Information Fig. S1 for "Optimization of Crop Rotation Calendar to Maximize System-Level Productivity under Climate Change"

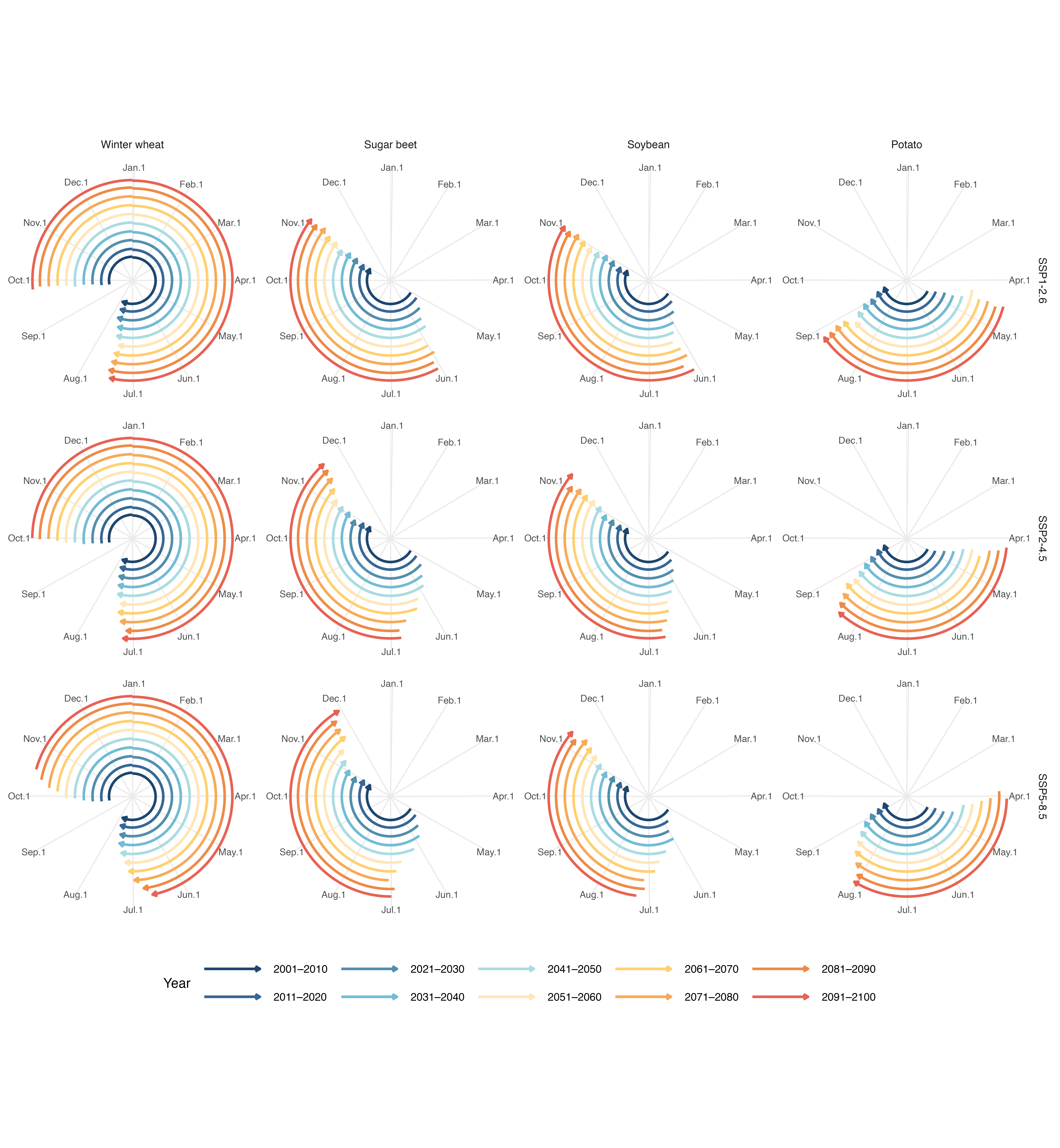

Fig. S1: Crop calendars of four rotation crops identified by the optimization algorithm for winter wheat-sugar beet-soybean-potato rotation under different SSP scenarios and time periods. The start and end points of arrows indicate mean sowing and harvesting dates, respectively.
