## Supporting Information Table S1 for "Optimization of Crop Rotation Calendar to Maximize System-Level Productivity under Climate Change"

Table S1: Numbers of available cropping options per year under different SSP scenarios and times periods. 10-year mean values and their minimum to maximum ranges are shown in parentheses.

| SSP | Year | Winter wheat | Sugar beet | Soybean | Potato |
| --- | --- | --- | --- | --- | --- |
| SSP1-2.6 | 2001–2010 | 389.3 (327.4-440.7) | 1481.4 (1096.2-1732.3) | 2036.8 (1438.6-2346.6) | 534.3 (483.2-559.9) |
| SSP1-2.6 | 2011–2020 | 402.1 (352.9-444.7) | 1788.4 (1536.7-1967.9) | 2632.3 (2202.0-3312.9) | 592.9 (558.5-631.5) |
| SSP1-2.6 | 2021–2030 | 415.0 (379.1-473.5) | 1726.5 (1563.7-1900.5) | 2693.1 (2436.5-3018.3) | 595.6 (546.4-626.9) |
| SSP1-2.6 | 2031–2040 | 406.9 (372.1-485.2) | 1651.1 (1267.0-2009.1) | 2575.1 (2056.5-2814.8) | 587.3 (493.1-634.4) |
| SSP1-2.6 | 2041–2050 | 382.3 (364.9-400.6) | 1653.5 (1526.4-1803.5) | 2566.3 (2176.8-2879.0) | 586.7 (555.7-625.0) |
| SSP1-2.6 | 2051–2060 | 366.0 (300.3-388.6) | 1641.9 (1378.7-1886.8) | 2466.2 (2130.8-2827.2) | 594.3 (549.5-629.1) |
| SSP1-2.6 | 2061–2070 | 409.7 (364.8-465.7) | 1774.5 (1514.1-2198.4) | 2626.3 (2480.2-2850.5) | 618.4 (556.9-646.1) |
| SSP1-2.6 | 2071–2080 | 417.3 (383.6-491.6) | 1879.6 (1514.4-2447.9) | 2793.2 (2590.4-3077.2) | 629.8 (586.9-671.4) |
| SSP1-2.6 | 2081–2090 | 418.9 (377.1-464.3) | 1756.2 (1570.8-1915.7) | 2744.7 (2411.5-3337.3) | 599.9 (553.2-654.7) |
| SSP1-2.6 | 2091–2100 | 403.6 (359.2-442.1) | 1656.9 (1419.8-2171.9) | 2554.1 (2137.8-2984.0) | 595.2 (549.1-648.8) |
| SSP2-4.5 | 2001–2010 | 389.3 (327.4-440.7) | 1481.4 (1096.2-1732.3) | 2036.8 (1438.6-2346.6) | 534.3 (483.2-559.9) |
| SSP2-4.5 | 2011–2020 | 396.7 (361.6-423.5) | 1809.8 (1691.6-1896.4) | 2544.4 (2124.6-2857.8) | 589.4 (564.3-609.3) |
| SSP2-4.5 | 2021–2030 | 388.4 (347.6-416.2) | 1605.8 (1356.5-1942.4) | 2521.5 (1715.9-3071.3) | 567.8 (525.9-612.2) |
| SSP2-4.5 | 2031–2040 | 388.0 (302.8-427.5) | 1567.4 (1215.9-1839.7) | 2587.4 (1834.7-2924.5) | 597.4 (523.0-631.0) |
| SSP2-4.5 | 2041–2050 | 404.4 (339.1-471.6) | 1805.7 (1347.3-2513.1) | 2729.8 (2254.6-3171.1) | 628.3 (569.8-689.4) |
| SSP2-4.5 | 2051–2060 | 397.9 (333.1-444.3) | 1742.1 (1432.7-1833.7) | 2620.9 (2243.3-2859.0) | 601.0 (555.7-641.3) |
| SSP2-4.5 | 2061–2070 | 405.7 (364.9-480.8) | 1818.1 (1335.0-2177.6) | 2832.4 (2298.9-3824.3) | 626.0 (553.7-686.0) |
| SSP2-4.5 | 2071–2080 | 409.1 (376.4-450.0) | 1762.9 (1508.3-2074.5) | 2593.4 (2380.9-2904.9) | 623.5 (587.0-643.6) |
| SSP2-4.5 | 2081–2090 | 386.6 (331.2-429.7) | 1745.8 (1421.5-2183.2) | 2489.9 (2217.2-2930.5) | 611.5 (568.4-662.1) |
| SSP2-4.5 | 2091–2100 | 397.2 (367.9-412.4) | 1748.8 (1527.8-2006.8) | 2540.4 (2397.2-2750.2) | 634.0 (585.4-676.7) |
| SSP5-8.5 | 2001–2010 | 389.3 (327.4-440.7) | 1481.4 (1096.2-1732.3) | 2036.8 (1438.6-2346.6) | 534.3 (483.2-559.9) |
| SSP5-8.5 | 2011–2020 | 388.4 (357.4-412.8) | 1781.8 (1630.2-1969.7) | 2562.2 (1947.5-2891.4) | 581.3 (551.5-614.4) |
| SSP5-8.5 | 2021–2030 | 390.0 (341.0-443.9) | 1647.2 (1494.3-1875.9) | 2449.1 (2034.5-2591.2) | 565.9 (534.0-608.0) |
| SSP5-8.5 | 2031–2040 | 395.4 (351.1-436.5) | 1700.2 (1549.2-1964.0) | 2526.2 (2336.5-3012.1) | 591.8 (560.2-630.5) |
| SSP5-8.5 | 2041–2050 | 391.8 (344.9-422.9) | 1660.9 (1431.7-1867.4) | 2647.1 (2432.3-2765.9) | 620.6 (573.1-659.4) |
| SSP5-8.5 | 2051–2060 | 387.3 (330.8-466.3) | 1760.3 (1595.3-2018.4) | 2567.0 (2172.2-3116.4) | 618.2 (591.8-630.8) |
| SSP5-8.5 | 2061–2070 | 429.6 (381.1-483.4) | 1934.1 (1773.5-2095.1) | 2586.2 (2324.6-2814.6) | 656.7 (628.8-678.1) |
| SSP5-8.5 | 2071–2080 | 375.9 (356.1-398.8) | 1959.6 (1611.8-2344.7) | 2400.4 (2196.1-2564.2) | 631.5 (592.1-681.1) |
| SSP5-8.5 | 2081–2090 | 363.0 (308.3-403.7) | 2067.8 (1728.1-2595.1) | 2288.0 (2064.9-2495.7) | 619.4 (533.3-682.7) |
| SSP5-8.5 | 2091–2100 | 327.7 (255.3-441.3) | 2137.8 (1755.4-2581.8) | 2269.5 (1927.6-2729.2) | 628.2 (482.4-694.4) |
