## Supporting Information Script S1 for "Optimization of Crop Rotation Calendar to Maximize System-Level Productivity under Climate Change": ScriptS1.R

```
library(tidyverse)

cropping_options <- read_rds("SupData/DataS1.rds")

optimize <-
  function(opt, crop_order, margin = c(10, 365), verbose = FALSE) {

    # fixed initial cropping option
    initial <-
      opt |>
      dplyr::filter(crop == crop_order[1],
                    year(sow) == min(year(sow))) |>
      dplyr::slice_max(output, with_ties = FALSE)

    # remove options to include only those after the initial harvest
    opt <-
      opt |>
      dplyr::filter(.data$sow >= initial$harvest[1]) |>
      dplyr::arrange(harvest) |>
      dplyr::bind_rows(initial, added_here = _)

    n <- nrow(opt)
    num_crop <- length(crop_order)

    # initialize dynamic programming tables
    # score[i,j], maximum cumulative output at i-th options with j-th crop
    # fin[i,j]: harvest date of the optimal path at i-th options with j-th crop
    score <- fin <- matrix(0, ncol = n, nrow = num_crop)

    # selected[[j]][[i]]: selected options at i-th option with j-th crop
    selected <- purrr::map(1:num_crop, ~ purrr::map(seq_len(n), ~ 0))

    # fill tables with the first option
    score[crop_order[1],1] <- opt$output[1]
    fin[,1] <- opt$harvest[1]
    selected[[crop_order[1]]][[1]] <- 1

    # filling dynamic programming tables
    for (i in 2:n) {

      crop_i <- opt$crop[i]
      # determine the preceding crop in the rotation order
      crop_prec <- c(crop_order[num_crop], crop_order)[which(crop_order == crop_i)]

      score[-crop_i, i] <- score[-crop_i, i - 1]
      fin[-crop_i, i] <- fin[-crop_i, i - 1]
      for(j in 1:num_crop){
        if(crop_i != j){
          selected[[j]][[i]] <- selected[[j]][[i-1]]
        }
      }

      new_weight <- opt$output[i]

      # find acceptable preceding options:
      accepts <- which(dplyr::between(as.numeric(opt$sow[i]) - fin[crop_prec, 1:(i - 1)], margin[1], margin[2]))

      if (length(accepts) != 0) {
        # if acceptable options exist, find the one with maximum cumulative output
        # choose the earliest sowing date for ties
        optima <- min(accepts[which(score[crop_prec, accepts] == max(score[crop_prec, accepts]))])
        new_weight <- new_weight + score[crop_prec, optima]
        candidate <- c(selected[[crop_prec]][[optima]], i)

        if ((new_weight > score[crop_i, i-1]) & identical(candidate[1], 1)) {
          # update DP tables if new cumulative output is greater than previous
          fin[crop_i, i] <- opt$harvest[i]
          score[crop_i, i] <- new_weight
          selected[[crop_i]][[i]] <- candidate
        } else {
          # otherwise, keep previous optimal values
          fin[crop_i, i] <- fin[crop_i, i-1]
          score[crop_i, i] <- score[crop_i, i-1]
          selected[[crop_i]][[i]] <- selected[[crop_i]][[i-1]]
        }
      } else {
        # if no acceptable options, keep previous optimal values
        fin[crop_i, i] <- fin[crop_i, i-1]
        score[crop_i, i] <- score[crop_i, i-1]
        selected[[crop_i]][[i]] <- selected[[crop_i]][[i-1]]
      }
    }

    # determine the highest-output pathway at the final position
    pathway <- which(score[,n] == max(score))

    if(!verbose){
      return(opt[selected[[pathway]][[n]],])
    } else {
      return(list(result = opt[selected[[pathway]][[n]],], score, fin, selected))
    }
  }


# demonstration -----------------------------------------------------------

opt <- cropping_options |> dplyr::filter(SSP == "SSP5-8.5", GCM == "MI", year(sow) %in% 2050:2062)

# print(opt)
#
# # A tibble: 72,694 × 6
# SSP      GCM   crop  sow        harvest    output
# <chr>    <chr> <chr> <date>     <date>      <dbl>
# 1 SSP5-8.5 MI    wheat 2050-10-04 2051-07-16  5018.
# 2 SSP5-8.5 MI    wheat 2050-10-04 2051-07-17  5156.
# 3 SSP5-8.5 MI    wheat 2050-10-04 2051-07-18  5269.
# 4 SSP5-8.5 MI    wheat 2050-10-04 2051-07-19  5348.
# 5 SSP5-8.5 MI    wheat 2050-10-04 2051-07-20  5414.
# 6 SSP5-8.5 MI    wheat 2050-10-04 2051-07-21  5445.
# 7 SSP5-8.5 MI    wheat 2050-10-04 2051-07-22  5471.
# 8 SSP5-8.5 MI    wheat 2050-10-04 2051-07-23  5512.
# 9 SSP5-8.5 MI    wheat 2050-10-04 2051-07-24  5518.
# 10 SSP5-8.5 MI    wheat 2050-10-04 2051-07-25  5518.
# # ℹ 72,684 more rows
# # ℹ Use `print(n = ...)` to see more rows


# convert ‘crop’ from chr -> int
# crop = 1: beet, crop = 2: potato, crop = 3: soya, crop = 4: wheat
opt <- opt |> mutate(crop = as.integer(as.factor(crop)))

identified <- optimize(opt, crop_order = c(4, 1, 3, 2))

print(identified)
# # A tibble: 13 × 6
#    SSP      GCM    crop sow        harvest    output
#    <chr>    <chr> <int> <date>     <date>      <dbl>
#  1 SSP5-8.5 MI        4 2050-10-17 2051-07-31  6073.
#  2 SSP5-8.5 MI        1 2052-05-01 2052-10-02  9689.
#  3 SSP5-8.5 MI        3 2053-06-09 2053-11-13  6735.
#  4 SSP5-8.5 MI        2 2054-04-18 2054-08-31 13503.
#  5 SSP5-8.5 MI        4 2054-09-27 2055-07-21  6250.
#  6 SSP5-8.5 MI        1 2056-05-22 2056-11-08 14240.
#  7 SSP5-8.5 MI        3 2057-06-14 2057-11-09  5930.
#  8 SSP5-8.5 MI        2 2058-03-28 2058-08-08 12184.
#  9 SSP5-8.5 MI        4 2058-10-04 2059-07-07  7904.
# 10 SSP5-8.5 MI        1 2060-06-07 2060-11-24 10373.
# 11 SSP5-8.5 MI        3 2061-06-23 2061-11-19  5503.
# 12 SSP5-8.5 MI        2 2062-04-18 2062-08-30 12580.
# 13 SSP5-8.5 MI        4 2062-10-02 2063-07-10  6784.
```
